# Using Drosophila to model a variant of unknown significance in the human cardiogenic gene *Nkx2.5*

**DOI:** 10.1101/2023.06.28.546937

**Authors:** TyAnna L. Lovato, Brenna Blotz, Cayleen Bileckyj, Christopher A. Johnston, Richard M. Cripps

## Abstract

Sequencing of human genome samples has unearthed genetic variants for which functional testing is necessary to validate their clinical significance. We used the Drosophila system to analyze a variant of unknown significance in the human congenital heart disease gene, *Nkx2*.*5*. We generated an R321N allele of the *Nkx2*.*5* ortholog *tinman* (*tin*) to model a human K158N variant and tested its function in vitro and in vivo. The R321N Tin isoform bound poorly to DNA in vitro and was deficient in activating a Tin-dependent enhancer in tissue culture. Mutant Tin also showed a significantly reduced interaction with a Drosophila Tbox cardiac factor named Dorsocross1. We generated a *tin*^*R321N*^ allele using CRISPR/Cas9, for which homozygotes were viable and had normal heart specification, but showed defects in the differentiation of the adult heart that were exacerbated by further loss of *tin* function. We conclude that the human K158N mutation is likely pathogenic through causing both a deficiency in DNA binding and a reduced ability to interact with a cardiac cofactor, and that cardiac defects might arise later in development or adult life.

## Introduction

Approximately 1% of children born in the US suffer some form of congenital heart condition that can lead to significant mortality or morbidity (Hoffman & Kaplan 2002; Reller et al 2008). Even when these conditions are surgically repaired, the patient can suffer long-term effects upon cardiac performance and overall health. Clearly, a better understanding of the processes that are affected as the heart is developing will provide insight into mechanisms for understanding and treating congenital heart disease. As a result, there has been intense focus upon defining the regulatory network controlling heart development (Waardenberg et al 2014).

Research over the past 30 years has determined that heart formation arises through the action of a conserved transcriptional and signaling network, that promotes heart cell specification and development (for reviews, see for example Bodmer & Venkatesh 1998; Cripps & Olson 2002; Xia et al 2020). During this time, the value of engaging a diverse group of model organisms to generate mechanistic understanding of cardiac development and disease has been exemplified by studies of the *tinman/Nkx2*.*5* genes. The gene *tinman* (*tin*) was first discovered in Drosophila (Kim & Nirenberg 1989) and shown to be both expressed in and required for the formation of the Drosophila heart (Bodmer 1993; Azpiazu & Frasch 1993). Subsequently, the mammalian ortholog of *tin* was identified as *Nkx2*.*5* (Lints et al 1993; Komuro & Izumo 1993) and shown to be required for normal heart development in mice (Lyons et al 1995). Shortly afterwards, mutations in human *Nkx2*.*5* were associated with congenital heart defects in three separate human pedigrees (Schott et al 1998). There have since been numerous examples demonstrating that conserved processes control heart development across the animal kingdom, and that mutations in these conserved genes impact cardiogenesis across multiple organisms (McCulley and Black, 2012).

Through this research, it has become apparent that a significant proportion of instances of human congenital heart defects arise from mutations in cardiac transcription factors. With the advent of whole-exome or whole-genome sequencing, a trove of natural variants in many transcription factor genes has been identified and deposited in databases such as ClinVar (https://www.ncbi.nlm.nih.gov/clinvar/). Many of these mutations are known to be pathogenic based upon clear association with clinical outcomes, as is the case with several *Nkx2*.*5* variants (see for example Schott et al 1998; McElhinney et al 2003). Furthermore, studies have sought to define the functional significance of human *Nkx2*.*5* mutations using in vitro and tissue culture studies (Kasahara & Benson, 2004), to assess the impact of mutations in the context of cardiomyocytes differentiating in vitro (Bouveret et al 2015), and some disease alleles have been modeled in mice (Costa et al 2013; Furtado et al 2017).

More broadly, studies in the mouse system have been successful in defining the developmental impacts of a number of mutations known to cause congenital heart disease, affecting several known cardiogenic genes (Majumdar et al 2021). Most of these studies have focused upon known pathogenic mutations, whereas mutations with unknown molecular or clinical effects have been under-studied. Mutations in this latter category should be a focus of study, however, since it is important to know if those changes require more detailed analysis to determine if they might contribute to human disease. Moreover, there are an increasing number of human variants of unknown impact (Telenti et al 2016), and the pace of identification of new variants has outstripped the ability to functionally or practically test them in vertebrate systems to determine if they might have clinical significance.

Given the conserved nature of heart specification and development across higher animals, one approach to efficiently test the functional significance of human variants is to model them in genetically amenable animals. This allows whole-organism studies to be carried out for a particular mutation in a more high-throughput manner, and can provide insight into the potential clinical relevance of a newly-discovered human variant. Here, we test this approach by modeling in Drosophila *tin* a basic (Arginine in Drosophila, Lysine in human) to polar Asparagine mis-sense mutation identified in *Nkx2*.*5*, for which the clinical significance is unknown. We use biochemical tests in vitro, and a genome edited in vivo allele, to demonstrate that this mutation attenuates protein function and results in reproducible defects in adult heart structure. We conclude that this mutation is likely to be pathogenic in humans, and propose that modeling human mutations in Drosophila can be an effective way to assess the functional significance of a variant.

## Materials and Methods

### Drosophila stocks and crosses

Drosophila were maintained at 25ºC on Jazz Mix medium (Genesee Scientific) for all experiments. Genetic nomenclature is as described at FlyBase.org (Gramates et al 2022). Embryos used for CRISPR were from flies carrying an X-linked *vas-Cas9* construct. The stock was obtained from the Bloomington Drosophila Stock Center (BDSC), #51323.

### CRISPR/Cas9 genome editing

CRISPR/Cas9 genome editing was used to generate an R321N allele of the endogenous tin gene. sgRNAs targeting *tin* close to codon R321 (protospacer sequence 5’-CGACTCAAAAAGTATCTGAC) and *ebony* (5’-GCCACAATTGTCGATCGTCA; used as a co-CRISPR target; Kane et al 2017) were ordered from Horizon Discovery Biosciences. A single-stranded donor DNA, comprising the intended nucleotide substitutions plus 50nt on each side, was also ordered from Horizon Discovery Biosciences. These reagents were sent to Rainbow Transgenic Flies Inc. for injection. G0 adults were crossed to a multiple balancer line containing *TM2* and *TM6* on the third chromosome. G1 generation *ebony* flies were then crossed to a deficiency line for *tin* balanced over TM3, *Sb* to test for lethality and to generate a stable stock. Genomic DNA from homozygous progeny was subjected to PCR and DNA sequencing to detect the presence of the intended point mutation.

### Protein purification

The Tin homeodomain (HD; codons 301-360) was cloned into the pMAL vector, containing an N-terminal maltose binding protein (MBP) tag, using 5’-NdeI and 3’-XhoI restriction sites. The R321N missense mutation was introduced using site directed mutagenesis PCR. The Doc1 coding sequence was cloned into the pGEX vector, containing an N-terminal GST tag, using 5’-BamHI and 3’-XhoI restrictions sites. All constructs were sequence confirmed prior to protein expression.

All proteins were expressed in BL21(DE3) *E. coli* under induction of isopropyl β-d-1-thiogalactopyranoside (IPTG) and grown in standard Luria–Bertani broth supplemented with 100μg/ml ampicillin. Transformed cells were grown at 37°C to an OD600 ∼0.6 and induced with 0.2mM IPTG overnight at 18°C. Cells were harvested by centrifugation (5000×*g* for 10 min), and bacterial pellets were resuspended in lysis buffer and flash-frozen in liquid nitrogen. Cells were lysed using a Branson digital sonifier and clarified by centrifugation (12,000×*g* for 30 min).

For MBP-tagged Tin, cells were lysed in lysis buffer (50 mM Tris pH 8.0, 500mM NaCl, 2mM DTT) and coupled to amylose resin for 3 h at 4°C. Following extensive washing, proteins were eluted with elution buffer (50mM Tris pH8, 30mM NaCl, 2mM DTT, 50mM maltose). Final purification was carried out using an S200-Sephadex size exclusion column equilibrated in storage buffer (20mM Tris pH 8.0, 200mM NaCl, 2mM DTT). Protein was concentrated and stored flash frozen at −80°C until subsequent use. For GST-Doc1, cells were lysed in lysis buffer as above and clarified lysate was stored flash frozen at −80°C until subsequent use.

### Protein methods

Electrophoretic mobilty shift assays were performed using standard procedures. The DNA binding buffer contained 40mM KCl, 15mM HEPES pH 7.9, 1mM EDTA, 0.5mM DTT, 5% Glycerol, and 4μg Poly-dIdC. The binding site was the Tin1 sequence from the *Multiplexin* gene (Lovato et al, in preparation), sequence 5’-GGACTCTTTGGCTGAAGTGTCCGGTGAAA, ordered from Sigma (Tin consensus binding sequence underlined). Equal quantities (1.5μg) of purified wild-type Tin-HD and Tin^R321N^-HD were used in each reaction.

Direct Doc1-Tin binding was assessed using standard GST pulldown assays. First, equivalent amounts of GST-Doc1 were coupled to glutathione agarose resin in separate reactions for 30 min at room temperature and extensively washed with PBS to removed unbound protein. Protein-bound resin was then resuspended in wash buffer (20 mM Tris pH8, 150 mM NaCl, 1 mM DTT, 0.2% Triton X-100) and incubated in the absence or presence of Tin (WT or R321N). GST alone was used as a negative control. For saturation binding experiments (see Figure 3), Tin proteins were added at increasing concentrations between 0-500nM in respective reactions. Reactions were then washed four times with wash buffer. Following the final wash, samples were resuspended in reducing Laemmli buffer and heated at 80°C prior to being resolved by SDS-PAGE. Due to the similar molecular weights of GST-Doc1 and MBP-Tin, MBP-Tin was detected by western blot using a rabbit anti-MBP antibody (GeneTex) and visualized using a Bio-RAD ChemiDoc imaging system. After detection, blots were stained with Ponceau red to visualize GST-Doc1 and ensure equivalent loading and transfer. Quantification of bound MBP-Tin was performed using ImageJ.

### Tissue culture

Co-transfection assays were carried out essentially as described by Kelly Tanaka et al (2008). The wild-type *tin* expression plasmid was pPAc-tinman (Lovato et al 2015). The *tin*^*R321N*^ mutant expression plasmid was generated using the Q5 site-directed mutagenesis kit (New England Biolabs). The reporter plasmid was the Mp3D cardiac enhancer from the *Multiplexin* (*Mp*) gene (Lovato et al, in preparation) fused to *lacZ* in the parent vector pDONR lacZ attB (Bryantsev et al 2012). Reporter expression was assessed using the mammalian β-galactosidase assay kit (Pierce Technology, Thermo Scientific). Reporter expression in the presence of transcription factors was normalized to reporter expression in the presence of empty expression vector. Ten replicates were carried out, and the reporter activities analyzed using One-way ANOVA and post-hoc Tukey tests.

### Generation of anti-Tin antiserum

The coding sequence of *tin* was cloned into the pEXP1-DEST vector (ThermoFisher V96001) utilizing Gateway Technology (ThermoFisher 12536017). A positive clone was transformed into BL21 (DE3) competent cells (ThermoFisher C600003), which was induced for protein purification using Talon Metal Affinity Resin (Takara 635501). Two New Zealand white rabbits were injected at ten sites with 500μl of Freund’s Complete Adjuvant (Sigma-Aldrich F5881) containing 500μg of purified Tin protein. A small blood draw was carried out 4 weeks later to check progress and a booster injection of 100μg of Tin protein in 200 μl of Freund’s Incomplete Adjuvant (Sigma-Aldrich 344291) was carried out at four sites. Two weeks later 50 ml of blood was drawn to check titer, after which a second booster was administered a week later. Four weeks later the final blood draw was obtained. Serum was collected by allowing the blood to coagulate at room temperature for 30 minutes and then centrifuged at 2000xg. The supernatant was transferred to a clean tube and analyzed by immunohistochemistry using a concentration of 1:1000. Approximately 50ml of blood was collected for each blood draw, and the rabbits were adopted. The protocol was IACUC approved (protocol number 18-200741-MC) prior to commencing the work.

### Immunofluorescence and microscopy

Embryos were harvested, fixed and stained as described by Patel (1994). Primary antibodies were mouse anti-Seven-up (1:100, Developmental Studies Hybridoma Bank); rabbit anti-Odd (1:1000, provided by Dr. James Skeath), and guinea pig anti-H15 (also known as Nmr1; 1: 2000, Dr. James Skeath). At least ten embryos of each genotype were analyzed, and representative images are shown.

Preparation and staining of larval pelts and of adult abdomens to visualize the heart was performed as described by Molina and Cripps (2001). The primary antibodies used were anti-ß-Integrin (CF.6G11; 1:25 dilution, DSHB) and anti-Alpha-actinin (1:2000, DSHB). The secondary antibody was Alexa 555 goat anti-mouse (1:2000, Invitrogen), combined with Alexa 488-Phalloidin (1:500, Invitrogen). At least five animals of each genotype were analyzed, and representative images are shown.

All samples were imaged using an Olympus FluoView 3000 confocal microscope. Images were assembled in Photoshop, and any changes to brightness, levels and contrast were applied equally to control and mutant samples.

To quantify the irregularity in Integrin stain for control and mutant samples, confocal images were imported into ImageJ (imagej.nih.gov). To measure the length of integrin stain, for each sample a freehand line was drawn over the integrin line. This was masked and then skeletonized, which enabled the determination of the length od the skeleton in pixels. In parallel, a separate line was drawn to best fit the ventral midline of the heart. The irregularity of the Integrin stain was calculated by dividing the length of the integrin line (in pixels) by the length of the best-fit line (in pixels) to create the Ratio of curve/straight as shown in Figure 5C. At least 15 samples were analyzed in this way for each genotype. Control and mutant samples were compared using Welch’s two-sample t-test in RStudio Cloud (now called Posit Cloud).

## Results

### In vitro analysis of the R321N mutation

The K158N variant of human *Nkx2*.*5* was submitted to ClinVar in 2017 (accession VCV000373685.1) and classified to be of unknown significance. Residue K158 of Nkx2.5 lies at the end of the first alpha-helix of the homeodomain (Pradhan et al 2012) and is one of three amino acids shown to directly contact Tbx5 in a ternary structure with DNA (Luna-Zurita et al 2016). These observations suggest that the K158N mutation might have clinical impact, but no functional testing is available to corroborate this hypothesis.

In the orthologous Drosophila gene, *tinman* (*tin*), residue R321 of Tin corresponds to K158 in Nkx2.5, which conserves the basic nature of the amino acid (Figure 1A). To determine if the Nkx2.5 variant might affect protein function, we modeled the basic-to-Asn mutation in *tin* and assessed its activity in vitro and in vivo. We first generated native protein corresponding to the homeodomains of wild-type (WT) and R321N mutant Tin fused to maltose binding protein (MBP) and purified these to near-homogeneity (Figure 1B). Next, we used each protein in an electrophoretic mobility shift assay to determine if the proteins could bind to a known Tin binding site from the *Multiplexin* (*Mp)* gene. Whereas the WT MBP-Tin homeodomain bound robustly to the radioactive probe (Figure 3C, lane 2), MBP-Tin homeodomain containing the R321N change reproducibly only bound weakly to the target sequence (Figure 3C, lane 3), indicating that the point mutation affects protein-DNA interaction.

**Figure 1:**
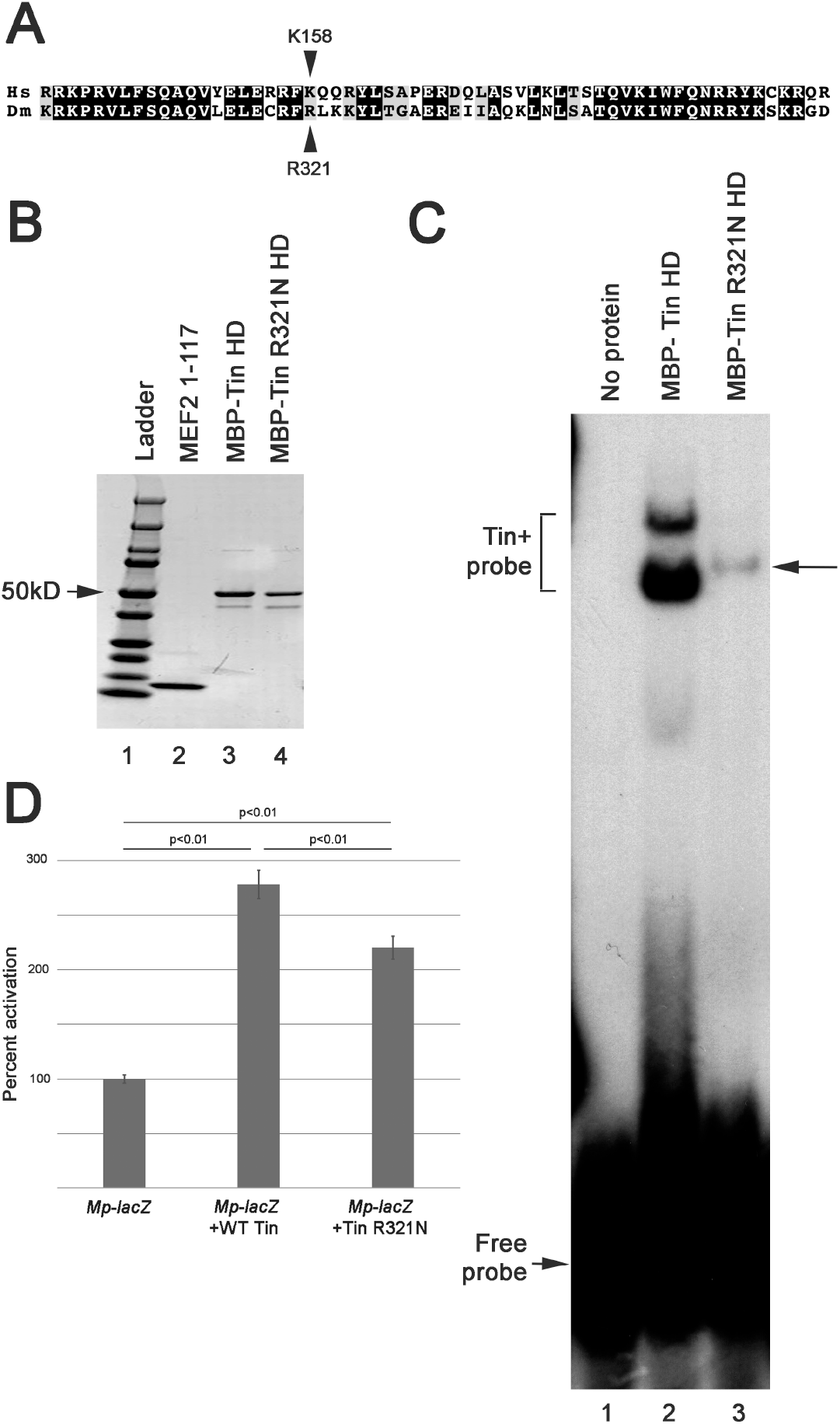
Functional analysis of Tin^R321N^ in vitro. A: Comparison of the homeodomain sequences of human (Hs) Nkx2.5 and *Drosophila melanogaster* (Dm) Tinman. The location of the point mutation being analyzed is indicated. B: Purified native proteins used for DNA binding assays. Equivalent amounts of wild-type MBP-Tin (lane 3) and mutated MBP-Tin (lane 4) were purified. The purified MEF2 was used for a different project. C: Electrophoretic mobility shift assay testing the ability of wild-type MBP-Tin homeodomain (MBP-Tin HD) and MBP-Tin^R321N^ HD to bind to a consensus Tin binding site in the *Multiplexin* (*Mp*) gene. Note that a Tin+Probe complex is formed in both the wild-type and mutant lanes (lanes 2 and 3), but that the intensity of the shifted probe is strongly reduced in the mutant, despite adding the same amount of purified protein. This electrophoretic mobility shift image is a representative image from two replicates. D: Co-transfection assays assessing the ability of wild-type and mutant Tin to activate an enhancer-*lacZ* reporter from the *Mp* gene. While both Tin isoforms can activate the reporter above background levels, the TinR321N shows a significantly reduced activation ability compared to wild-type. Data are from ten replicates. One-way ANOVA of the three sample sets determined significant differences between the treatments (p<0.0001), and post-hoc Tukey tests were applied to each pairwise comparison.

We further tested the impact of the R321N mutation in transient co-transfection experiments. Here, we transfected into Drosophila S2 tissue culture cells expression plasmids expressing WT or R321N-mutant full-length Tin, plus a reporter plasmid comprising an *Mp* cardiac enhancer fused to the *lacZ* gene. Transfected cells were lysed and assessed for the accumulation of ß-Galactosidase (ßGal) expressed from the reporter, normalized to ßGal expression in the presence of empty expression plasmid. We found that WT Tin activated the reporter significantly, whereas the R321N isoform activated reporter expression more weakly.

Nevertheless, the reduced activation from the mutant construct was still significantly greater than baseline activation (Figure 1D)

Taken together, these results indicate that the R321N mutation attenuates Tin function, but that some significant activity still remains in the mutant isoform.

### Interaction of Tin with the Tbx ortholog Dorsocross 1

The crystal structure of a complex between DNA, Nkx2.5 and Tbx5 was recently described (Luna-Zurita et al 2016; Pradhan et al 2016), and interestingly one of the three residues of Nkx2.5 that interacts directly with Tbx5 is K158. Moreover, combined mutation of K158 plus the two other interacting residues reduced the interaction between Nkx2.5 and Tbx5 (Luna-Zurita et al 2016). In Drosophila, while no physical interaction of Tin with any Tbx proteins has been described, a genetic interaction between *tin* and three *Tbx6* orthologs (named *Dorsocross* (*Doc*) *1-3*) has been documented. Specifically, loss of function for *tin* increases the severity of cardiac patterning defects arising from *Doc* mutations (Reim & Frasch 2006). We therefore sought to determine if this genetic interaction is also reflected at the protein level. In addition, given the strong conservation in Tbox sequences between paralogs, we determined if the strength of any interaction between Tin and a Doc protein was affected by the R321N mutation.

We first determined if a physical interaction between Tin and Doc could be documented. We selected Doc1 to test, since the T-box DNA binding domain in this isoform is identical to that of Doc2, and these two isoforms are most similar to human Tbx6. We performed pulldown experiments using purified MBP-Tin wild-type and R321N prey protein and tested their interaction with GST-Doc1 bait protein immobilized on glutathione agarose. We found that whereas MBP alone had negligible binding (Figure 2, lane 7), significant binding of MBP-Tin with GST-Doc1 was evident (Figure 2, lane 8). This binding was absent when MBP-Tin was tested with GST alone as a control (Figure 2, lane 4). These data demonstrate for the first time that Tin can physically interact with a T-box protein. The MBP-Tin R321N mutant also bound to GST-Doc1 (Figure 2, lane 9), but this binding appeared to be diminished compared to wild-type protein.

**Figure 2:**
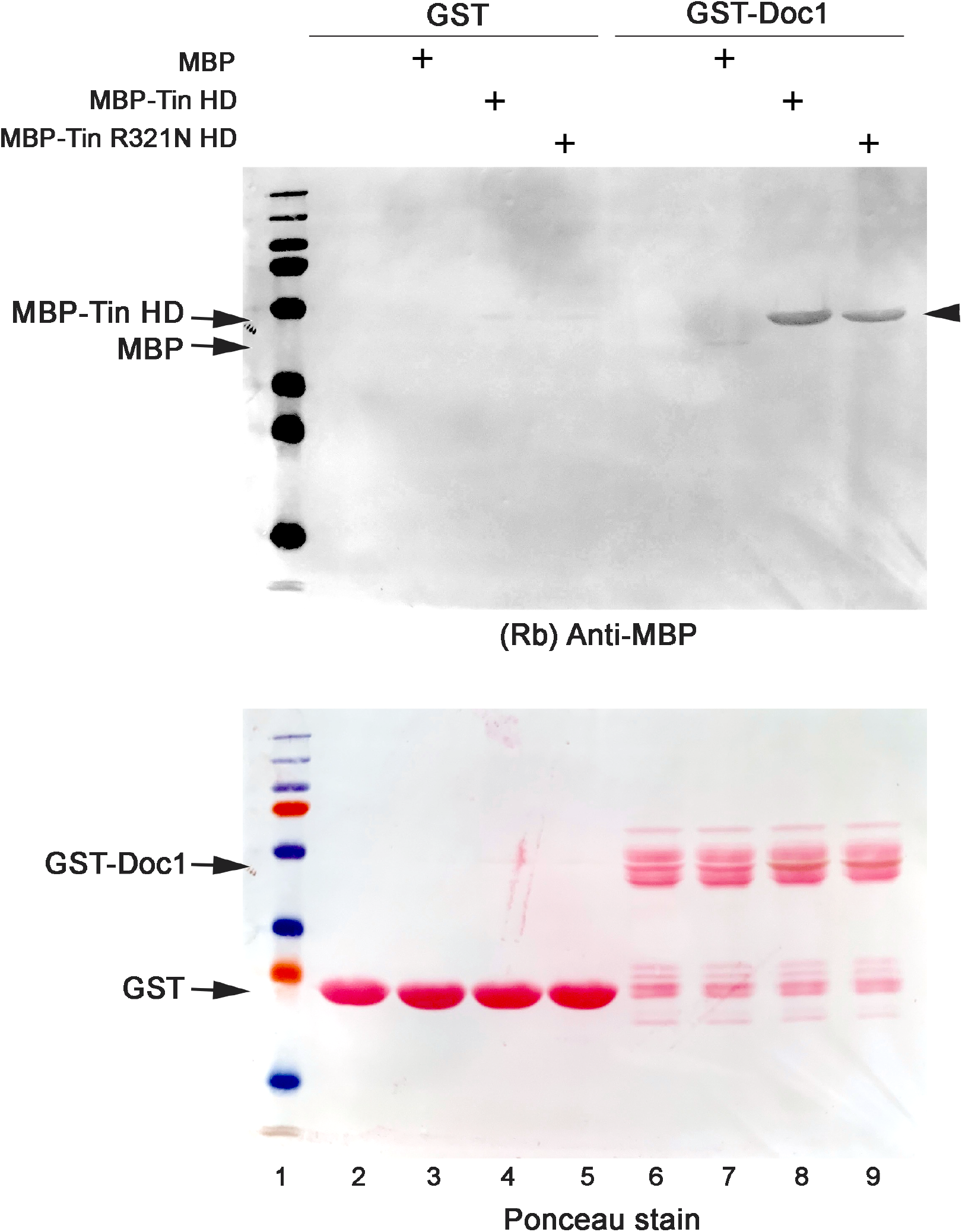
The Tin homeodomain interacts directly with Doc1. GST alone or fused to Doc1 (GST-Doc1) were immobilized as solid phase bait on glutathione agarose, washed extensively, and subsequently incubated without or with 10 μM MBP alone or MBP-Tin (wild-type or R321N mutant) prey proteins as indicated. Following incubation, reactions were extensively washed and resolved by SDS-PAGE. Due to the overlapping molecular weights of GST-Doc1 and MBP-Tin, we visualized bound MBP-Tin in western blots using an anti-MBP antibody (top panel). Ponceau red staining of the blot was done following western blot developing to ensure equal GST and GST-Doc1 loading and transfer. Binding to GST was minimal for all MBP proteins. GST-Doc1 did not show a significant interaction with MBP alone but did bind both MBP-Tin wild-type and R321N at this single concentration. Binding to the R321N appeared reduced compared to wild-type.

These initial interaction studies examined only a single concentration of MBP-Tin proteins binding to GST-Doc1. To determine the affinity of these interactions and better assess if this interaction is affected by the R321N substitution, we next performed saturation binding experiments by examining binding across a lower range of MBP-Tin protein concentrations. As shown in Figure 3, the wild-type MBP-Tin bound robustly across nanomolar concentrations (Figure 3A) with kinetics consistent with a one-site saturation binding (Figure 3B). In contrast, MBP-Tin R321N was significantly impaired at these lower concentrations, revealing a severe impact of this mutation on Doc1 binding. These data demonstrate that Tin binds Doc1 with a high affinity and that the R321N mutant dramatically impairs this interaction.

**Figure 3:**
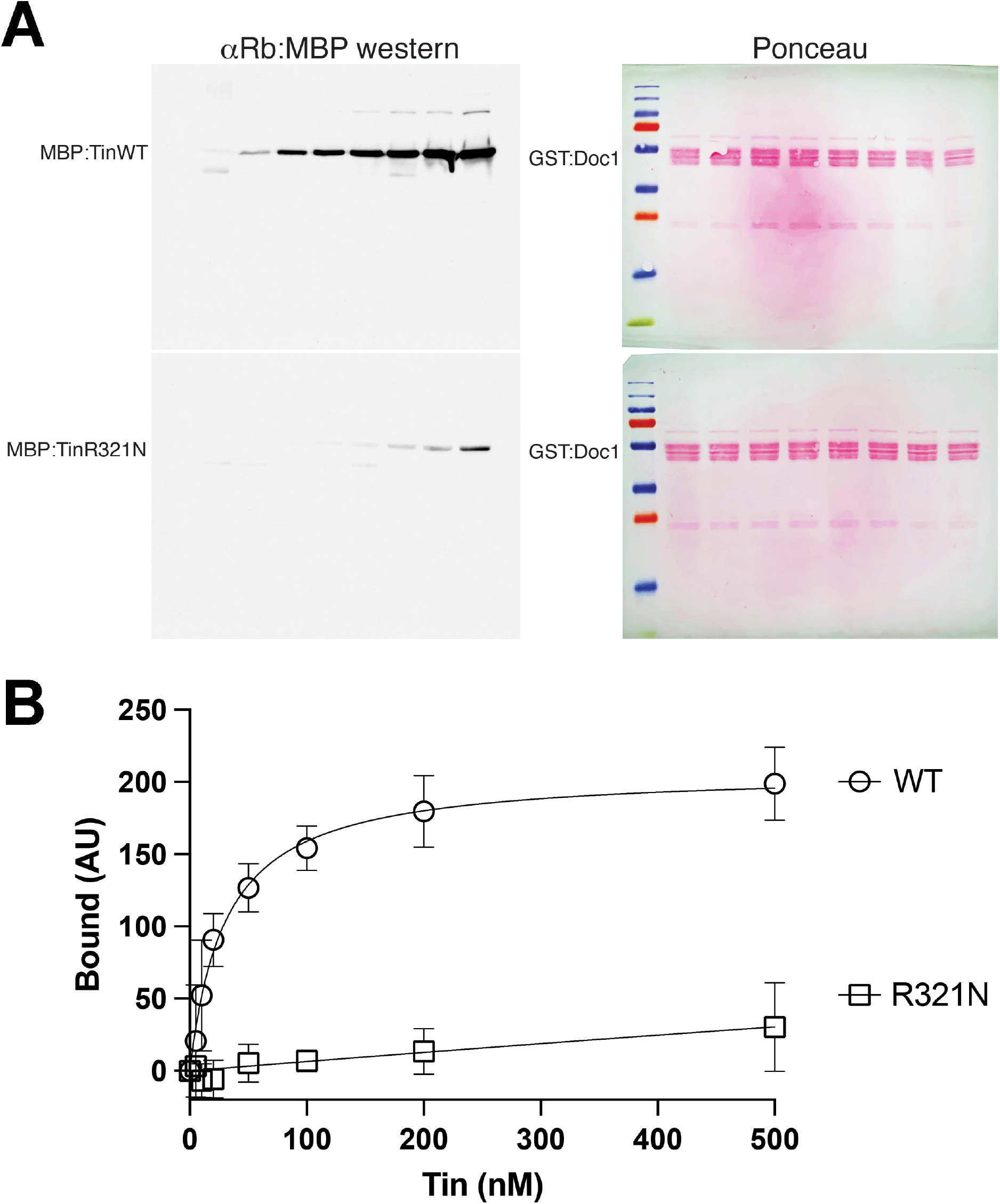
The TinR321N isoform interacts with Doc1, but with a significantly reduced affinity. A: Equal amounts of GST-Doc1 were immobilized on glutathione agarose, washed extensively, and subsequently incubated in the presence of increasing concentrations (0 – 500nM) of MBP-Tin wild-type or R321N mutant. Following incubation, reactions were extensively washed and resolved by SDS-PAGE. Bound MBP-Tin proteins were visualized by western blot analysis using an anti-MBP antibody (Left panels); GST-Doc1 was visualized using Ponceau red staining of blots (Right panels). Wild-type MBP-Tin (Top panel) bound robustly across the concentration range tested, whereas the R321N mutant (Bottom panel) was significantly impaired in GST-Doc1 binding. B: Saturation binding curves of the average ± standard deviation from five independent experiments demonstrates the effects of R321N mutation on Doc1 binding. Bound MBP-Tin proteins were analyzed in ImageJ and normalized to the total GST-Doc1 for respective samples.

Taken together, our in vitro and tissue culture studies indicate that the R231N mutation significantly impacts several aspects of Tin function, including DNA binding, transactivation of target genes, and interaction with an essential cardiac co-factor.

### Generation and analysis of a tin^R321N^ mutant in vivo

We next sought to determine if this mutation affects Tin function in the context of the intact animal. We used CRISPR/Cas9 genome editing to generate a 5-’CGA to 5’-AAC codon change and isolated one line that showed this alteration with no other changes to the *tin* coding sequence (Figure 4A). Interestingly, these flies were homozygous viable and fertile, and could be maintained as an homozygous stock.

**Figure 4:**
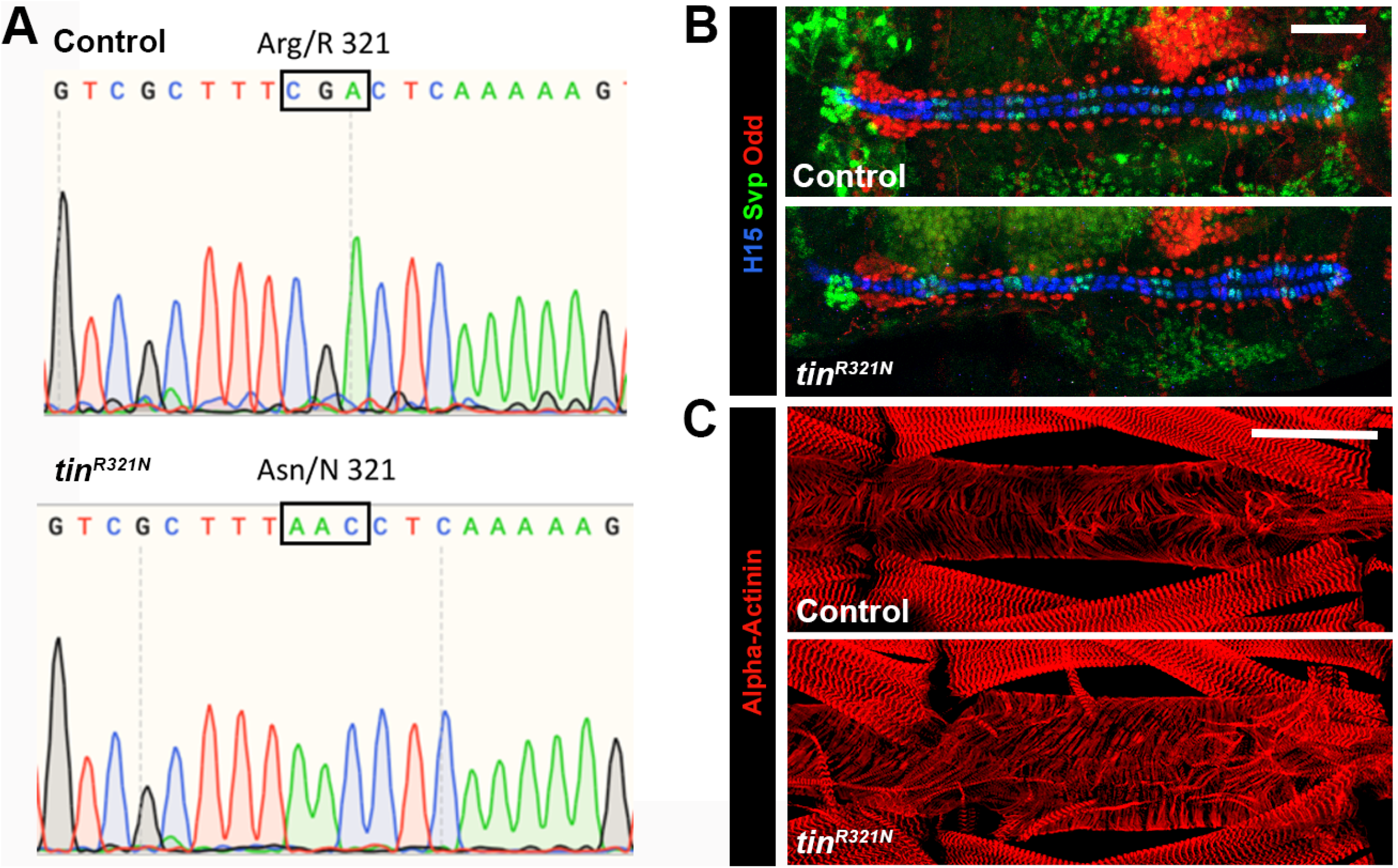
Generation and analysis of a *tin*^*R321N*^ allele in vivo. A: Sequence reads of wild-type *tin* (top) and the *tin*^*R321N*^ allele (bottom), from PCR products generated from homozygous adults. The Arg codon (5’-CGA) is altered to Asn (5’-AAC), and no other changes to the tin coding sequence were observed. B: There were no observable changes in heart cell patterning of *tin*^*R321N*^ homozygotes compared to *vasa-Cas9* controls. Cardial cells are labelled with anti-H15 (Blue) and anti-Seven-up (Green). A subset of pericardial cells is labelled with anti-Odd (Red). Bar, 50μm. C: Larval hearts from the A6 abdominal segment of *vas-Cas9* (Control) and *tin*^*R321N*^ homozygous larvae, labeled with anti-Alpha-Actinin (Red) to visualize cardiac myofibrils. No consistent differences in heart structure were observed between control and mutant. Bar, 50μm.

To determine if the R321N mutation affected Tin function in vivo, we assessed heart formation in wild-type and homozygous-mutant embryos. In wild-type animals, the mature embryonic heart at stage 16 comprises two parallel rows of cardial cells forming a cardiac tube, flanked by a number of pericardial cells.

Within these two cell populations there is significant molecular diversity: the mature cardial cells express either *tin* or the orphan nuclear receptor gene *seven-up* (*svp*); and the pericardial cells express a number of different transcription factors, with a subset of the cells expression the C2H2 transcription factor gene, *odd-skipped* (*odd*) (Figure 4B, top). In *tin*^*R321N*^ homozygotes, we did not observe any consistent differences in cardial and pericardial cell patterning between control and mutant animals, suggesting that any effect of the point mutation upon *tin* function was not sufficient to impact heart cell specification nor patterning (Figure 4B, bottom).

We next determined if the point mutation of *tin* affected cardiac growth. During larval development, the cardiac tube generated in the embryo increases significantly in size, and then during pupal development metamorphoses into the adult heart (Molina and Cripps, 2001). When we studied the heart of fully grown third-instar larvae, we did not observe any differences in organization of the myofibrils between control and *tin*^*R321N*^ homozygotes (Figure 4C).

By contrast, when we studied adult heart structure, we observed defects in the arrangement of the adult heart cells in mutant animals. In wild-type, the persistent larval cardiac tube increases in size and undergoes extensive myofibrillogenesis to form a muscular tube (Molina and Cripps, 2001). Along the ventral side of this tube are arrayed a series of longitudinal muscle fibers (Figure 5A). The arrangements of individual heart tube cells can be visualized through staining for accumulation of ß-Integrin, that is enriched at the junctions of each cardial cell and is particularly visible along the ventral midline of the tube. In wild-type animals, this line of Integrin staining was relatively straight (Figure 5A), indicating that contralateral cells meet at approximately the same midline location. However, in *tin*^*R321N*^ homozygotes, the line was far more irregular: in all 17 samples analyzed this line was continuous but irregular, and frequently deviated significantly from the midline (Figure 5B).

**Figure 5:**
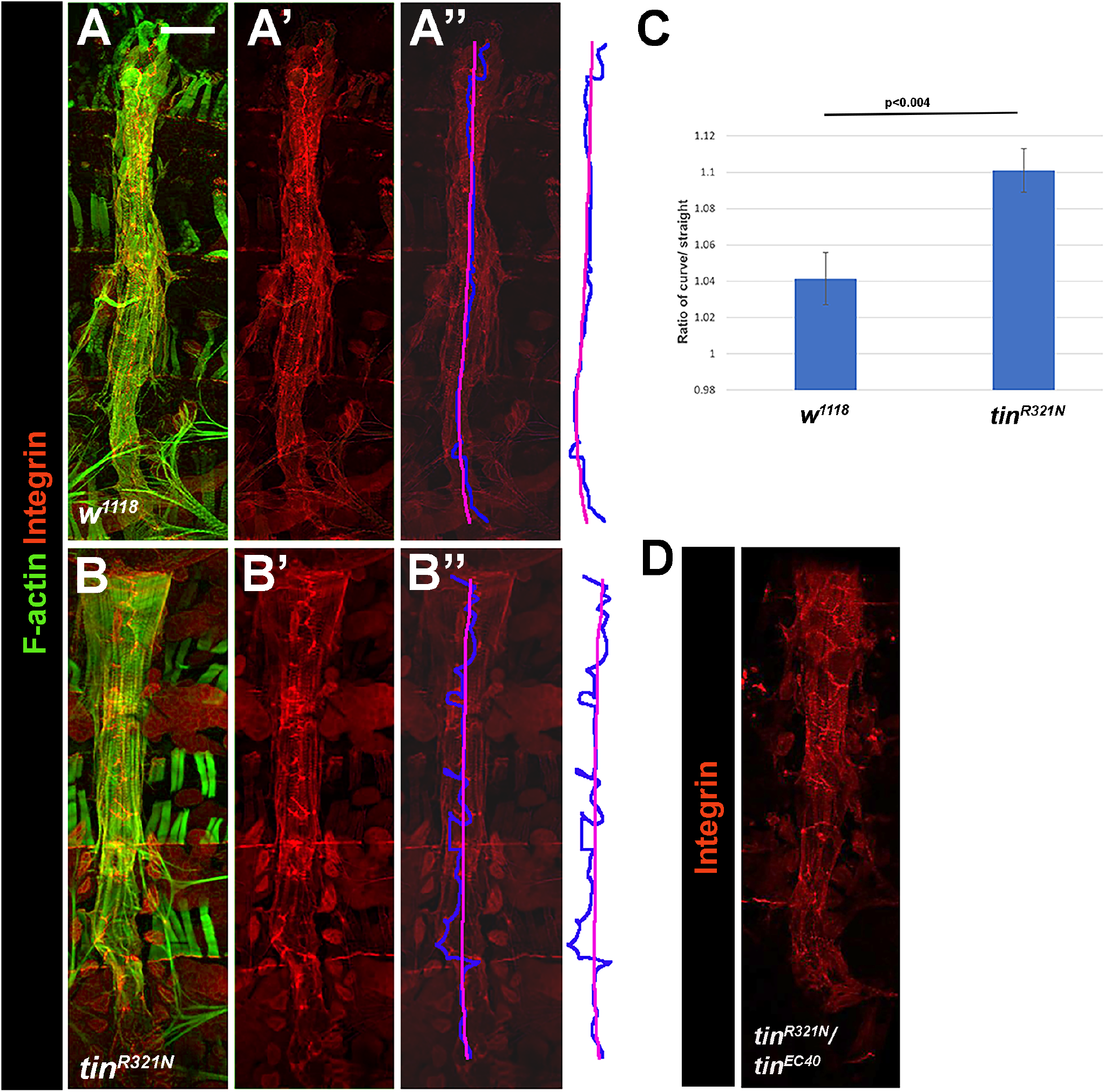
The *tin*^*R321N*^ adults have defects in adult heart cell patterning. A,B: Whole-mount preparations of *w*^*1118*^ (Control) and *tin*^*R321N*^ homozygous adult hearts, stained for accumulation of F-actin (Green) and ß-PS Integrin (Red). While the overall structure of the heart did not differ between genotypes based upon the F-actin accumulation, accumulation of Integrin was irregular in the mutants (B’) compared to control (A’). In controls, a single prominent line of Integrin stain was observed along the ventral midline of the heart, traced by a Blue line in A’’. This line of stain was irregular in the *tin*^*R321N*^ homozygotes (B’’). To quantify the irregular nature of the line, the length of the Blue line was calculated relative to that of a single straight line along the ventral midline (Magenta in A’’ and B’’), to give a ratio of curve/straight. C: The calculated ratio for *w*^*1118*^ was significantly less than that of *tin*^*R321N*^, indicating that the line was more irregular in the mutants compared to controls. D: Combining tinR321N with the tin null allele tinEC40 to create tinR321N/tinEC40 trans-heterozygotes resulted in severe disruptions to heart cell patterning where the ventral line of Integrin stain was highly disrupted and could not be consistently raced as a single line from anterior to posterior. Bar, 50μm.

To quantify this phenotype, the “length” of the Integrin line in pixels was calculated using online software (see Materials and Methods). The length of the Integrin line was then compared to a best-fit line representing the least distance along the ventral midline of the heart between the start and stop point of the integrin line (Figure 5 A’’), to generate a curve/straight ratio. A perfectly straight line of Integris stain would score a ratio of 1.0, whereas a line that deviated from the midline would show a ratio greater than 1.0. We calculated these ratios for control and *tin*^*R321N*^ homozygotes, where we observed that the mutant ratio was significantly greater than that for the control (Figure 5C, p<0.004).

We also addressed whether the defects we observed here were due to the engineered mutation of *tin*, and not due to a second-site mutation on the same chromosome. To achieve this, we crossed *tin*^*R321N*^ adults to heterozygotes for a *tin* null allele, *tin*^*EC40*^ (Bodmer 1993), and analyzed adult heart structure in *tin*^*R321N*^*/tin*^*EC40*^ adults. Since the *tin* null allele would provide zero Tin function, we would predict a stronger phenotype in these trans-heterozygous animals than in the *tin*^*R321N*^ mutants.

In the *tin*^*R321N*^*/tin*^*EC40*^ adults, we observed severely fragmented Integrin staining in the majority (9/11) of animals (Figure 5D), which could not be quantified in the manner described above due to the breakage in the line of staining. This enhancement of the mutant phenotype when in trans with a *tin* null allele indicated that the defects we observed arose from deficiencies in Tin function.

## Discussion

In this paper, we test the feasibility of using Drosophila as a model to understand the potential impact of a human variant of unknown significance in the congenital heart disease gene *Nkx2*.*5*. Modeling Nkx2.5^K158N^ as Tin^R321N^ generated a polypeptide that in vitro showed a strongly reduced ability to interact with DNA and reduced trans-activation ability in co-transfection experiments. Since the mis-sense mutation affects the DNA-binding homeodomain of Tin, our findings were consistent with reduced protein-DNA interactions for the mutant isoform. Based upon these effects of the mutation, we anticipated that modeling the same mutation in the endogenous *tin* gene in vivo would severely impact heart specification and differentiation. Instead, the *tin*^*R321N*^ homozygotes were viable and fertile, and showed only modest, albeit reproducible, defects in the patterning of cells in the adult heart.

We also demonstrate here for the first time a physical interaction between the T-box-encoded protein Doc1 and the cardiogenic factor Tin. While we only tested the ability of Doc1 to interact with Tin, we feel it is likely that all three Doc paralogs are capable of this interaction based upon their highly similar sequences and expression during heart development (Reim et al 2003; Reim & Frasch 2006). A physical interaction between Tin and Doc1 is also consistent with a genetic interaction between *tin* and the three paralogous *Doc* genes (Reim & Frasch 2006).

Our studies are also consistent with the documented interaction between mammalian Nkx2.5 and Tbx5 (Pradhan et al 2016; Luna-Zurita et al 2016). While Doc1 is not the closest Drosophila sequence ortholog of human Tbx5, the similar roles of Doc proteins in heart specification and the interaction of Doc1 with Tin/Nkx2.5 may indicate that these factors are functional orthologs. In support of this, one Nkx2.5 residue that lies at the interaction surface between Nkx2.5 and Tbx5 is K158 (Luna-Zurita et al 2016), that we show here to also be required for effective interaction of Tin with Doc1. These data suggest that there may be common mechanisms underlying the interactions of NK homeodomain proteins with Tbox proteins that could represent a mechanistic basis for multiple biological processes. In support of this, the NK homeodomain factor CEH-51 and the T-box factor Tbx-35 also genetically interact in the formation of the *C. elegans* pharynx (Briotman-Maduro et al 2009), and several examples exist of homeodomain proteins interacting with T-box proteins (Farin et al 2008; Nowotschin et al 2006; Leconte et al 2004; Boogard et al 2008).

At the molecular level, it is interesting to note that the R321N substitution affects both DNA binding and co-factor interaction, and these effects can provide insight into the molecular basis of the mutant phenotype and potential disease mechanisms. Homeodomains are known to fold into three alpha-helices, the third of which is the predominant region for interacting with DNA (Gehring et al 1990; Kissinger et al 1990). According to the Nkx2.5 crystal structure (Prabhan et al 2012), K158 is a surface-facing residue that lies at the C-terminal end of helix 1. Given the strong homology in homeodomain structure, R321 in Tin likely occupies the same structural location. Such a position on the “surface” of the homeodomain is consistent with a role in co-factor interaction with Doc, as we have confirmed here and as shown by Luna-Zurita et al (2016). However, how this mutation impacts DNA binding is less obvious. We note that the nearby Y162 residue in Nkx2.5 (Y325 in Tin) contacts DNA (Prabhan et al 2012), so it is possible that the R321N substitution impacts the ability of Y325 to connect properly with the DNA binding site. Alternatively, the R321 residue, given its location at the end of the first alpha-helix, might de-stabilize the structure of the helix, causing a reduced interaction of the homeodomain with DNA in vitro.

Our studies also raise a number of important considerations when the goal is to understand the potential clinical impact of uncharacterized mutations. First, there is a clear difference in the severity of the phenotypes when comparing assays performed in vitro versus in vivo. Based upon the DNA binding and co-transfection studies,Tin^R321N^ is clearly deficient compared to wild-type Tin, yet *tin*^*R321N*^ homozygotes show normal heart specification and early differentiation. Clearly, whatever functional deficiencies that were observed in vitro appear to be muted in vivo. This may be due to different biochemical environments, or due to the presence of co-factors that can stabilize some deficiencies in Tin function. Our results underline the important contributions that in vivo studies make to understanding the phenotypic severity for a given mutant allele.

Second, we show that there is an in vivo impact of the *tin*^*R321N*^ mutation and suggest strongly that in humans this mutation is likely to be pathogenic. That the phenotypes do not arise in flies until later in development, when the heart has undergone its maximum period of growth during metamorphosis, suggests that in humans this mutation might impact later stages of heart development.

Third, we note that the effects of the mutation upon adult flies are quite modest and are only exacerbated when heterozygous with a *tin* null allele. These observations underline the importance of accurate phenotypic analyses, that must be reproducible and quantifiable to confirm that subtle differences in heart development or function have occurred. Several groups have pioneered such assays in Drosophila that can be brought to bear on this issue (Cammarato et al 2015; Huang et al 2022).

Overall, this study demonstrates the feasibility of modeling human genetic variants in Drosophila, and opens a pathway for analysis of further variants of unknown significance using the Drosophila heart model.

## Acknowledgements

This work was supported by GM124498 awarded by the NIH to RMC, and Grant MCB-2205405 from the NSF to CAJ. CB is supported by a predoctoral fellowship from the American Heart Association. BB was supported by a Summer Undergraduate Research Program award from San Diego State University. We are grateful to Dr. James Skeath for the gifts of anti-H15 and anti-Odd.

